# Reorganization of Septin structures regulates early myogenesis

**DOI:** 10.1101/2023.08.24.554594

**Authors:** Vladimir Ugorets, Paul-Lennard Mendez, Dmitrii Zagrebin, Giulia Russo, Yannic Kerkhoff, Tim Herpelinck, Georgios Kotsaris, Jerome Jatzlau, Sigmar Stricker, Petra Knaus

## Abstract

Controlled myogenic differentiation is crucial for developmental formation, homeostatic maintenance and adult repair of skeletal muscle and relies on cell fate determinants in myogenic progenitors or resident stem cells. Proliferating muscle progenitors migrate, adopt spindle shape, align membranes and fuse into multinuclear syncytia. These processes are accompanied by cyto-architectural changes driven by rearranging of cytoskeletal components such as actin and microtubules. Here we highlight septins, the fourth component of the cytoskeleton, to represent an essential structural element of myoblasts. Specifically, Septin9 regulates myoblast differentiation during the early commitment process. Depletion of Septin9 in C2C12 cells and primary myoblasts led to a precocious switch from a proliferative towards a committed progenitor transcriptomic program. Additionally, we report Septin9 undergoing substantial reorganization and downregulation during myogenic differentiation. Together, we propose filamentous septin structures and their controlled reorganization in myoblasts to provide a key temporal regulation mechanism for the differentiation of myogenic progenitors.

## Introduction

Skeletal muscle is a dynamic tissue making up about 40% of the body and is essential for many vital functions^1,2^. The process of building muscle, i.e. myogenesis, is a multistage process where myogenic cells undergo a series of differentiation events from cycling progenitors (myoblasts) to fusion competent progenitors (myocytes) that ultimately fuse to form new myotubes, or enlarge existing myotubes by fusion^3,4^. These extensive morphological changes are accompanied by adaptations in cytoskeletal protein organization such as actin^5–7^ and tubulin^8,9^.

Septins constitute a family of GTP binding cytoskeletal proteins, that form non-polar filaments and just begin to emerge in the myogenic context^10^. They are divided into four subgroups (Septin2-Septin6-Septin7-Septin9) and vary mainly in N– and C-terminal extensions^11^. They assemble into palindromic hetero-octamers (henceforth protomer) with subunits from one of each of the four subgroups. Through lateral interactions, septins build higher-ordered structures such as bundles, rings and networks. The ability to bind either the plasma membrane, actin or microtubules defines their cellular functions as lateral diffusion barriers or compartmentalizing scaffolds^12–14^. Septins were reported in a wide range of cellular functions such as cytokinesis^15^, cell mobility^16^ and contractility^17^, plasma membrane rigidity^18,19^ and cell shape determination^20–23^, and mechanotransduction^24,25^. Here, we focus on Septin9, a centrally positioned paralog in the protomer^26^, the embryonic depletion of which is lethal^27^. The unique N-terminal extensions of Septin9 convey interaction with actin and microtubules^13^ and potentially link actin to the plasma membrane^28^. Expression, organization, and functions of septins during muscle formation are poorly understood. Human septin transcripts were detected in skeletal muscle and heart tissue^29^. Septin7 was required for zebrafish cardiac and somitic myofibril organization^30^, murine myoblast proliferation and skeletal muscle architecture and function^31^. Furthermore, Septin7 regulated the proliferative state of neuronal progenitor cells (NPCs) during cortical development^32^, and prevented premature aging of hematopoietic stem cells (HSCs) by compartimentalizing cytoplasmic polarity proteins^33^.

## Results and Discussion

In order to understand which septins are expressed in myogenic progenitors, we performed unsupervised trajectory analysis on single-nucleus RNA-sequencing (snRNA-seq) data of the developing musculature ranging from embryonic day 9.5 to 13.5^34^. Clusters annotated as muscle progenitors, myoblasts and myocytes were defined as myogenic lineage based on pseudotime expression of *Pax3*, *Pax7*, *Myod1* and *Myog* (Fig 1a,b and Fig 1a,b). This *in silico* approach revealed *Septin2*, *7*, *9* and *11* to be the highest expressed septins from every homology group. *Septin9* shows increased expression in myoblasts but is downregulated as these differentiate into myocytes (Fig 1c). From other expressed septins (1, 3, 4, 5, 6, 8 and 10) *Septin6* shows a strong downregulation in cells transitioning to myocytes while *Septin4*, virtually not expressed in myogenic progenitors, shows a strong upregulation in myoblasts followed by mild downregulation in myocytes (Fig EV1c). We confirmed the expression of septins (except *Septin3*, 12 and *14*, not detectable) during 7 days of *in vitro* myogenesis in C2C12 cells (Fig 1e,f and Fig EV2a-d) and primary myoblasts (5 days) (Fig EV2e-h), showing consistent downregulation of *Septin9* from 3 days on (Fig 1f and Fig EV2f). Additionally, the downregulation of *Septin6* and *7* and upregulation of *Septin4* were observed in C2C12 cells and primary myoblasts (Fig 1f and Fig EV2c-h). In line, protein level of Septin9 was downregulated during myogenic differentiation in C2C12 cells and primary myoblasts (Fig 1g and Fig EV2i). Based on these data, we confirmed the expression of most septins along myogenic differentiation and identified Septin2, 7, 9 and 11 as the highest expressed paralogs, constituting the core myoblast septin protomer (Fig 1d). Additionally, we report on potentially regulatory septin paralogs such as Septin4 and Septin6, which are described as highly regulated differentially expressed genes in C2C12 cells^35^. The sole upregulation of *Septin4* towards late commitment and fusion stage might represent a triggered response of the pro-apoptotic Septin4 splice variant Arts^36^ to an increased apoptotic resistance in mature myotubes^37–39^. Other less abundant myoblast septins such as Septin1, 5, 8 and 10 might be redundant, tune the septin filament composition or engage in core filament-independent functions. Together these results show that septins, especially Septin9, are expressed in myogenic progenitors and are mostly downregulated during myogenic differentiation.

**Figure 1.**
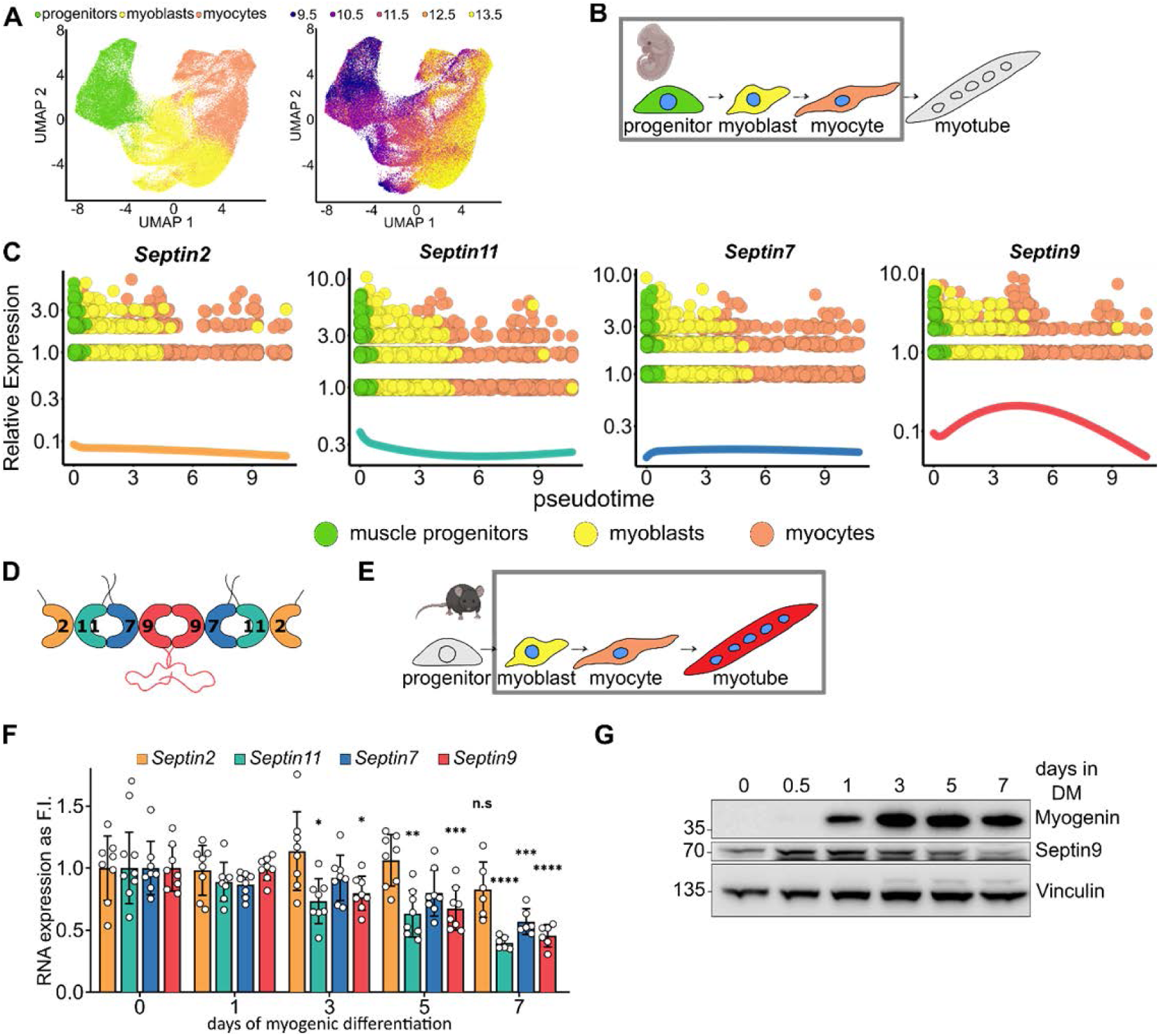
Expression of Septin9 during development and *in vitro* myogenesis. **a** UMAP visualization of the developing musculature ranging from embryonic day 9.5 to 13.5. Nuclei were isolated *in silico* from the mesenchymal cell trajectory, constructing a myocyte subtrajectory. Isolated cellular subtypes (left) and analyzed days of embryonic development (right) are color coded. **b** Schematic representation of murine embryonic myogenesis stages addressed in **a** and color coded according to cell states defined in **a**. **c** Pseudotime expression profiles of *Septin 2, 7, 9* and *11* in developing musculature. Spline curves representing the overall trend in gene expression along the differentiation trajectory are color coded according to the schematic in **d**. **d** Potential core myogenic septin protomer. **e** Schematic representation of murine *in vitro* myogenesis. **f** mRNA expression levels of *Septin2*, *7*, *9* and *11* during C2C12 differentiation. Septins are color coded according to the subfamily, as in **d**. **g** Protein levels of Septin2 and 9 during C2C12 differentiation. Data represent mean ± standard deviation (SD), *p<0.05, **p<0.01, ***p<0.001, ****p<0.0001, n.s – not significant from one way ANOVA followed by Dunnett’s multiple comparison test.

Septin9 plays a key role in dictating the subcellular localization of the septin cytoskeleton and in regulation of cytoarchitecture^28,40^. Therefore, we asked next if Septin9 downregulation merely removes a vestigial cytoskeletal component no longer required during late myogenesis or serves a potentially instructive purpose regulating the progression through myogenesis. Thus, we depleted Septin9 using an siRNA approach (Fig 2a) and observed a total loss of septin structures in proliferating myoblasts (Fig EV6c). We then performed transcriptomic analysis of Septin9-depleted differentiating C2C12 cells (after 12h of differentiation, Fig 2) and proliferating primary myoblasts (Fig EV3). Both data sets had a high overlap in expressed genes (Fig 2b). We compared the profiles of proliferating and committed muscle stem cells (MuSCs) defined by correlation with an external single-cell RNA-sequencing dataset (top 25 genes for each cluster^3^) in Septin9-deficient C2C12 cells and primary moyblasts (Fig 2c). Both Septin9-deficient C2C12 cells and primary myoblasts show upregulation of genes from the cluster “committed progenitors” and mild downregulation of genes associated with proliferation, suggesting precocious transition of the cycling progenitor population towards committed progenitors. Differential expression analysis (Fig 2d-g, Fig EV3b-d) revealed 201 differentially expressed genes (DEGs) in C2C12 cells, (65 up– and 136 downregulated) as compared to control cells transfected with non-targeting siRNA (Fig 2e). 213 DEGs were determined in Septin9-deficient primary myoblasts (168 up– and 45 downregulated) (Fig EV3c). Gene ontology annotation analysis of DEGs in Septin9-deficient C2C12 cells revealed association of upregulated genes with the terms ‘Myod1 targets’, ‘hallmark myogenesis’, ‘positive regulation of myoblast differentiation’, ‘positive regulation of skeletal muscle tissue development’ and ‘E box binding’ (Fig 2f). Downregulated DEGs are associated with ‘reduces cellular response to certain stimuli (TGFβ, TNFα, mTORC1)’, ‘cell cycle regulation (G2M transition, mitotic spindle)’ and ‘cell adhesion (cadherin binding, adherens junction assembly)’ (Fig 2f). Similar results were observed in Septin9-deficient proliferating myoblasts (Fig EV3d). 41 shared DEGs were found between C2C12 and primary myoblasts (Fig 2g). Most upregulated DEGs belong to a set of robustly regulated genes defined in a comparative study of nine RNA sequencing datasets from cycling or differentiating C2C12 cells ^35^. Some of those genes also belonged to the “hallmark myogenesis” GO term (marked red) such as *Myog*, *Actn3*, *Des* and *Cdkn1a (p21)*, pointing together towards upregulated myogenic differentiation (Fig 2g). Other upregulated and potentially Septin9 knockdown specific DEGs include (i) carbonic anhydrase 3 (Car3), an intracellular pH regulator and an early marker of myogenesis^41^ and (ii) matrix GLA protein (Mgp), an ECM protein inhibiting myostatin receptor binding^42^. Downregulated DEGs comprise a heterogenous selection of genes, such as (i) actin binding proteins such as *Dstn* and *Amotl2*, (ii) growth factors’ signal integrating SH2/SH3 containing adapter *Crkl*, (iii) soluble TGFβ ligand ActivinA (*Inhba*) and (iv) transcription factors DP1 (*Tfdp1*), Taz (*Wwtr1*) and beta-catenin (*Ctnnb1*) among others. Together, evaluation of DEGs points towards the premature switch from proliferating to differentiating myogenic progenitors in cells lacking Septin9.

**Figure 2.**
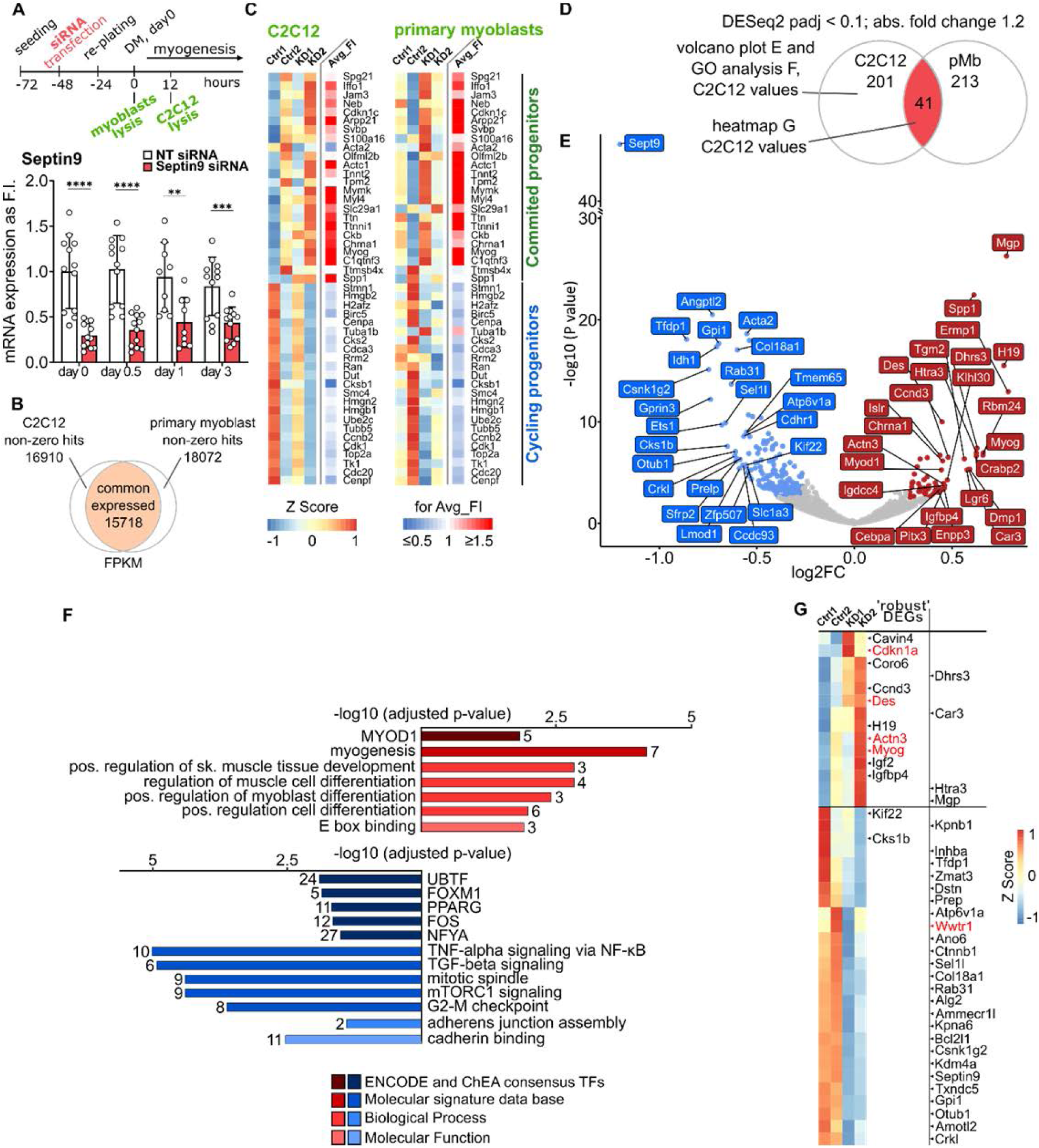
Septin9 depletion induces transition from cycling towards committed progenitors. **a** Schematic representation of the experimental setup. Total RNA was isolated from proliferating myoblasts 48 hours after Septin9 knockdown and from 12 hours differentiating C2C12 cells (60 hours after knockdown). Septin9 knockdown efficiency validation over 3 days of in vitro myogenesis via qRT-PCR in C2C12 cells. **b** Venn diagram comparing all expressed genes in C2C12 and primary myoblasts. **c**. Heat map depicting 47 genes in C2C12 (left panel) and in primary myoblasts (right panel) sorted into 2 clusters: ‘cycling progenitors’ and ‘committed progenitors’, annotation adapted from^3^ (except *Gm7325, 2810417h13rik, 2700094k13rik* not expressed in C2C12 and in primary myoblasts and excluded from the heatmap). Septin9 depletion leads to accelerated switch towards committed myogenic progenitor program, seen by general upregulation of genes in the cluster “committed progenitors” and mild downregulation of genes in the cluster “cycling progenitors”. Avg_FI represent the average between two experiments measuring KD to Ctrl ratio. **d** Venn diagram comparing DEGs calculated using DESeq2 (adjusted p value < 0.1; –0,322 ≤ log2FC ≤ 0,263). **e** Volcano plot of DE genes (adjusted p value <0.1; –0,322 ≤ log2FC ≤ 0,263) of C2C12 cells transfected with either non-targeting siRNA (control) or Septin9 siRNA after 12 hours of myogenic differentiation, 50 most regulated DEGs are highlighted. **f** GO enrichment analysis of genes upregulated (red) or downregulated (blue) in Septin9-deficient C2C12 cells. **g** Heatmap depicting 41 DEGs in C2C12 calculated using DESeq2 (adjusted p value < 0.1; –0,322 ≤ log2FC ≤ 0,263), highlighting in red genes from associated to the GO term “hallmark myogenesis”. Data represent mean ± standard deviation (SD), *p<0.05, **p<0.01, ***p<0.001, ****p<0.0001 from two-sided unpaired t test.

To understand whether an increased transcriptional signature translates into accelerated or premature myogenic differentiation in Septin9-deficient cells we analyzed the expression of early and late myogenic marker genes via qRT-PCR (Fig 3a-b). Upon robust reduction in Septin9 expression (reduction in proliferating cells to 30% and to 52% after 3 days of differentiation compared to control cells, Fig 2a) we show early expressed myogenin (*Myog*) up– and Taz (*Wwtr1*) down-regulation in proliferating and early differentiating C2C12 cells (Fig 3a). Later myogenic genes such as follistatin (*Fst*) and myosin heavy chain 2 and 8 (*Myh2* and *Myh8*), which are not expressed in cycling myoblasts and are induced during myogenesis, showed a significant upregulation after 72 hours of differentiation in C2C12 cells upon Septin9 knockdown (Fig 3b). We further confirmed the increased expression of Myog (after 24 hours) and Myh2 (after 72 hours) upon Septin9 depletion on the protein level (Fig 3c). In addition, we show that depletion of Septin9 leads to an early increase in myogenic differentiation index after 3 days of differentiation, assessed via quantification of myosin heavy chain 2 expression (Fig 3d,e). Furthermore, myoblasts lacking Septin9 exhibit no apparent early fusion defects, as addressed by myogenic fusion index (Fig 3f). In summary, loss of Septin9 in myoblasts associates with the early activation of myogenic target genes, resulting in premature differentiation.

**Figure 3.**
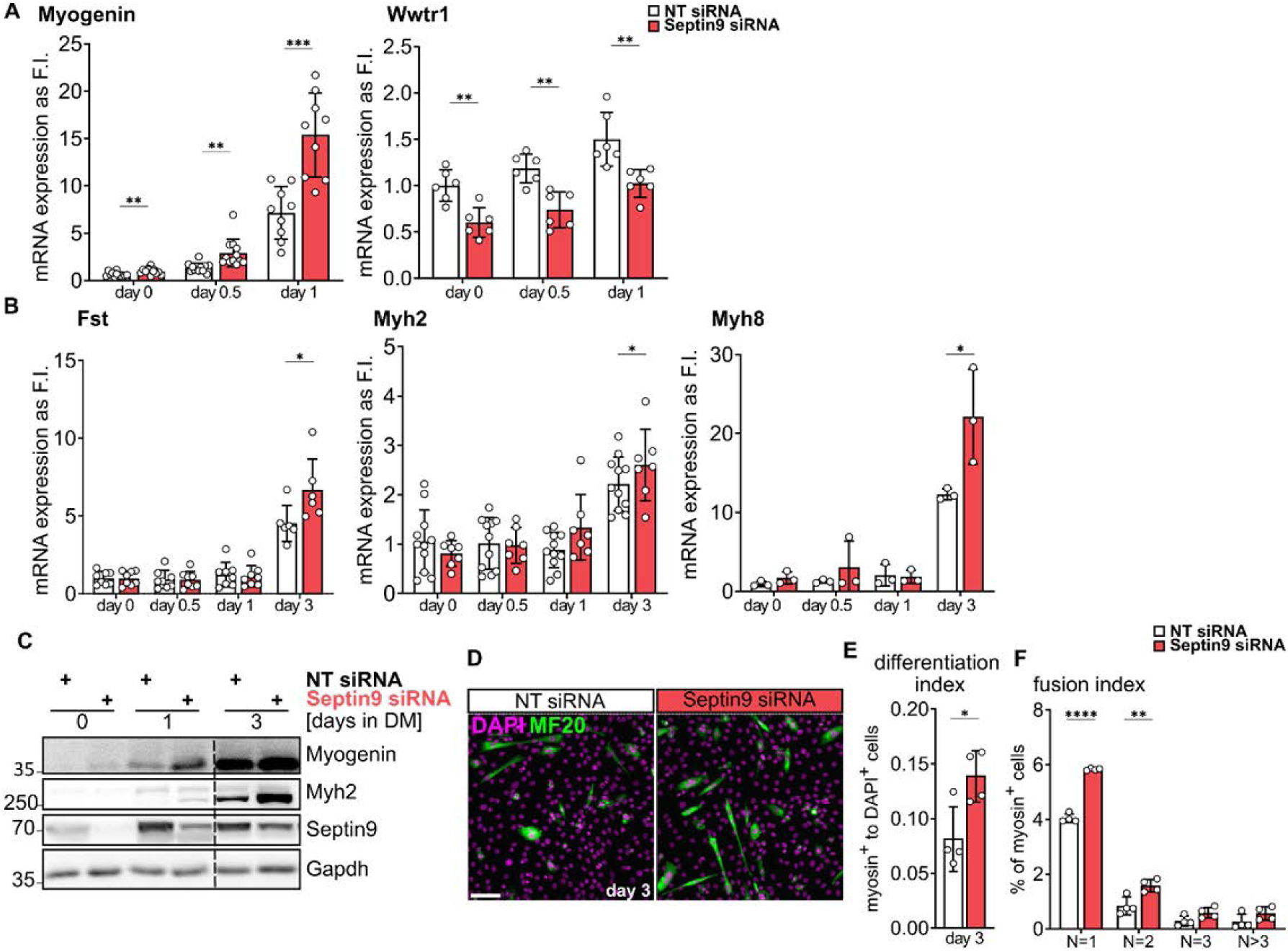
Septin9 depletion accelerates myogenic differentiation in C2C12 cells. **a**. qRT-PCR analysis of mRNA expression of DE expressed early and late (**b**) myogenic markers. **c** Western blot analysis of myogenic markers myogenin and myosin heavy chain 2 in Septin9 deficient myoblasts compared to control. **d** Immunofluorescence stainings of myosin heavy chain 2 (MF20) after 3 days of differentiation. **e** Myogenic differentiation index (MDI) and **f** myogenic fusion index (MFI) for cells in **d** respectively. N is the number of nuclei. Data represent mean ± standard deviation (SD), *p<0.05, **p<0.01, ***p<0.001, ****p<0.0001 from two-sided unpaired t test. Scale bar 50 µm.

Myogenic differentiation requires extensive reorganization of the myoblast cytoskeleton and acquisition of a migratory and fusion competent phenotype by myocytes^43,44^. Therefore, we next assessed the organization of Septin9-containing complexes during early myogenic differentiation. To follow the dynamic process of septin filament organization, localization and rearrangement during myogenesis, we generated a C-terminal meGFP-tagged Septin9 C2C12 line using CRISPR/Cas9 (Fig EV6a-b). In proliferating myoblasts, Septin9-GFP partially localized to perinuclear and ventral actin fibers, adjacent to focal adhesions (Fig 4a and Fig EV5a), but not to dorsal actin fibers (Fig 4a, inset ii), as observed before in other cell lines^28^. These data are in accordance with previous research showing Septin9 to directly cross-link actin filaments into bundles and promote maturation of nascent focal adhesions^45^. In contrast, myotubes show only remnants of filaments such as short rods, spirals and rings, sporadically distributed along the myotube, with only few perinuclear filaments remaining (Fig 4b,d). To shed light on septin reorganization during early myoblast differentiation and fusion, we performed live cell experiments with Septin9-GFP C2C12 cells. Septin reorganization close to the plasma membrane during early differentiation of proliferating myoblasts transitioning to myocytes were captured using total internal reflection (TIRF) microscopy. Imaging Septin9-GFP we observed structural changes events as early as 12 hours of differentiation (Fig 4c,e; Supplement Movie 1). During differentiation, septins successively became more dynamic and curved, partially losing their association to actin fibers (Fig 4c, Fig EV4). It is worth noting that Septin9 never localized *in vitro* to α-actinin in myoblasts (Fig EV4,5), which is involved into formation of sarcomeres and undergoes changes during myogenic differentiation^46^. During myocyte fusion assessed via confocal microscopy after 3 days of differentiation (Fig 4d), a different organization of septin structures for the invading versus the receiving myoblast was observed at the basal plane. The receiving myocyte had curvy perinuclear septin filaments whereas in the invading fusion-competent myocyte Septin9-GFP appeared reorganized intoshort rods and rings. These invading cells exhibited very mobile and largely septin-free lamellipodia that are known to be involved in the fusion process^47,48^, with Septin9 being present at the leading edge (Fig 4d, white arrow heads at 1:00 and 3:00). After membrane fusion, perinuclear filaments mixed and reorganized to receive a new fusion event (Fig 4d, at 7:30; Supplement Movie 2, 3). The nascent myotube (Fig 4d, at 14:00 and 15:30) exhibited septin filaments spanning across the nuclear region, while the rest of the cell contained rings and other filament remnants (Fig 4d red arrow heads at 14:00). The sarcoplasm, visible through the middle plane, appeared largely devoid of septins throughout the fusion process (Fig 4d, lower panels).From the live cell experiments, we conclude that actin-based septin filaments gradually disassemble from actin fibers during myogenesis, as myoblasts undergo morphological changes towards myocytes. Myocytes adopt a migratory character with largely septin-free lamellipodia and short curvy septins (Fig 4f). Freshly fused nascent myotubes have perinuclear dynamic fibers and only short remnants of filaments in the rest of the cell body. Considerably reduced association with actin during differentiation could be explained by shifts in affinity of septins towards microtubules or the plasma membrane^40,49–52^. Since septins colocalize only occasionally with microtubules in myoblasts and myotubes (Fig EV7), they might translocate to the plasma membrane, away from forming sarcomeric machinery and towards the increasingly dynamic cell surface of committed, fusion-competent progenitors^53,54^.

**Figure 4.**
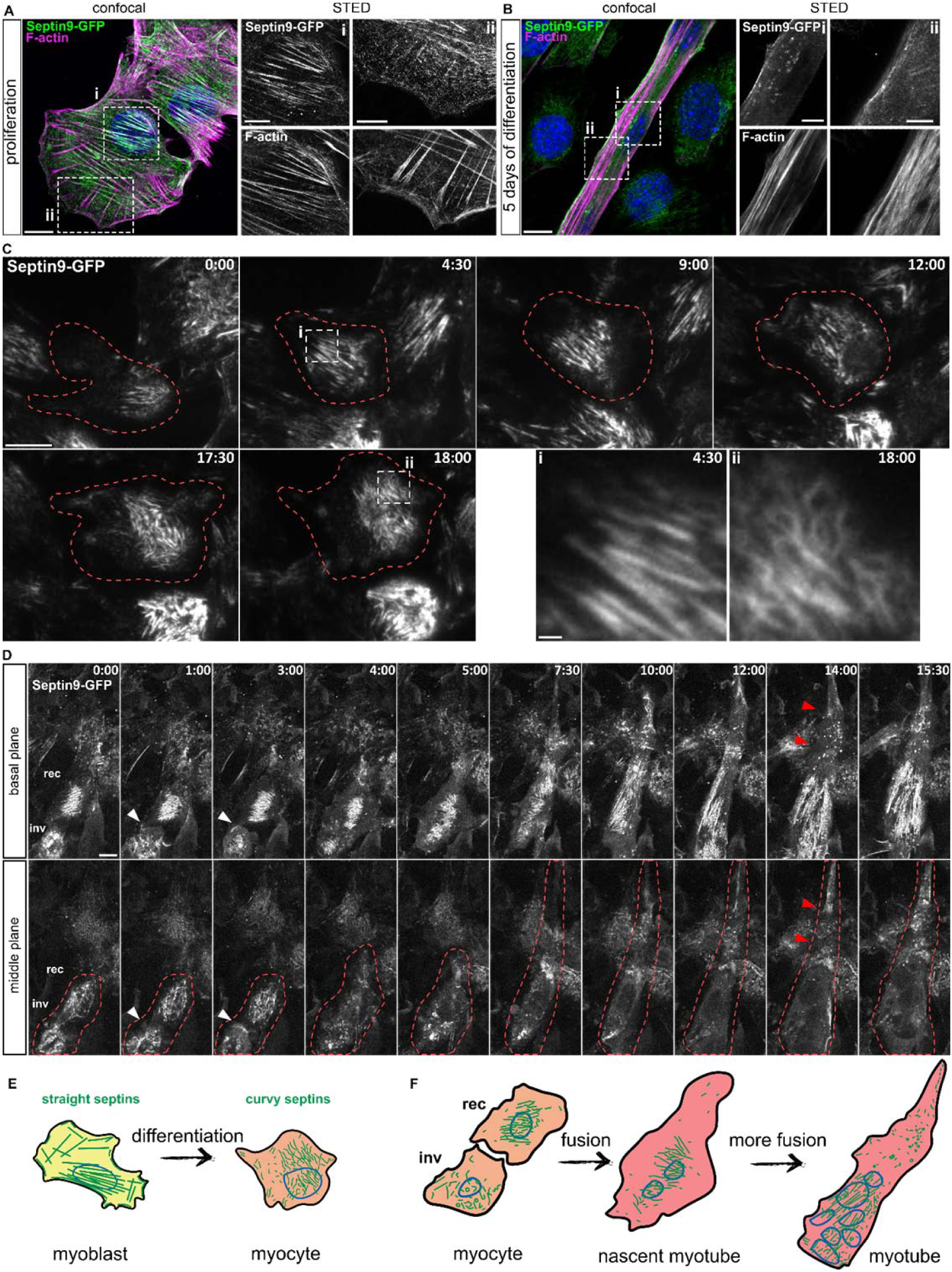
Septin9 reorganization during differentiation and fusion in Septin9-GFP C2C12 cells. **a**. Representative immunofluorescence micrographs showing septin filaments in proliferating Septin9-GFP C2C12 cells **a** and myotubes **b**. Merge in confocal, inlays in STED. **c** Representative snapshots from a live cell TIRF experiment using differentiating Septin9-GFP C2C12 cells. Acquisition started 12 hours after onset of myogenic differentiation, time in hours: minutes. Insets highlight changes in septin fiber morphology. **d** Representative snapshots from a live cell confocal experiment representing the basal and the middle plane of fusing Septin9-GFP C2C12 cells. Acquisition started 72 hours after onset of myogenic differentiation, time in hours:minutes. White arrow heads show Septin9-GFP at the leading edge of a lamellipodium. Red arrow heads show absence of Septin9-GFP in the distal part of a nascent myotube. **e** Schematic representation of a myoblast differentiating to a myocyte with according changes in septin morphology. **f** Schematic representation of a fusion event between to myocytes and the further maturation of the nascent myotube with according changes in septin morphology at several representative time points. Scale bar 10µm, insets 5 µm (a-b) and 1µm (c). inv-invading cells, rec-receiving cell.

**Figure 5.**
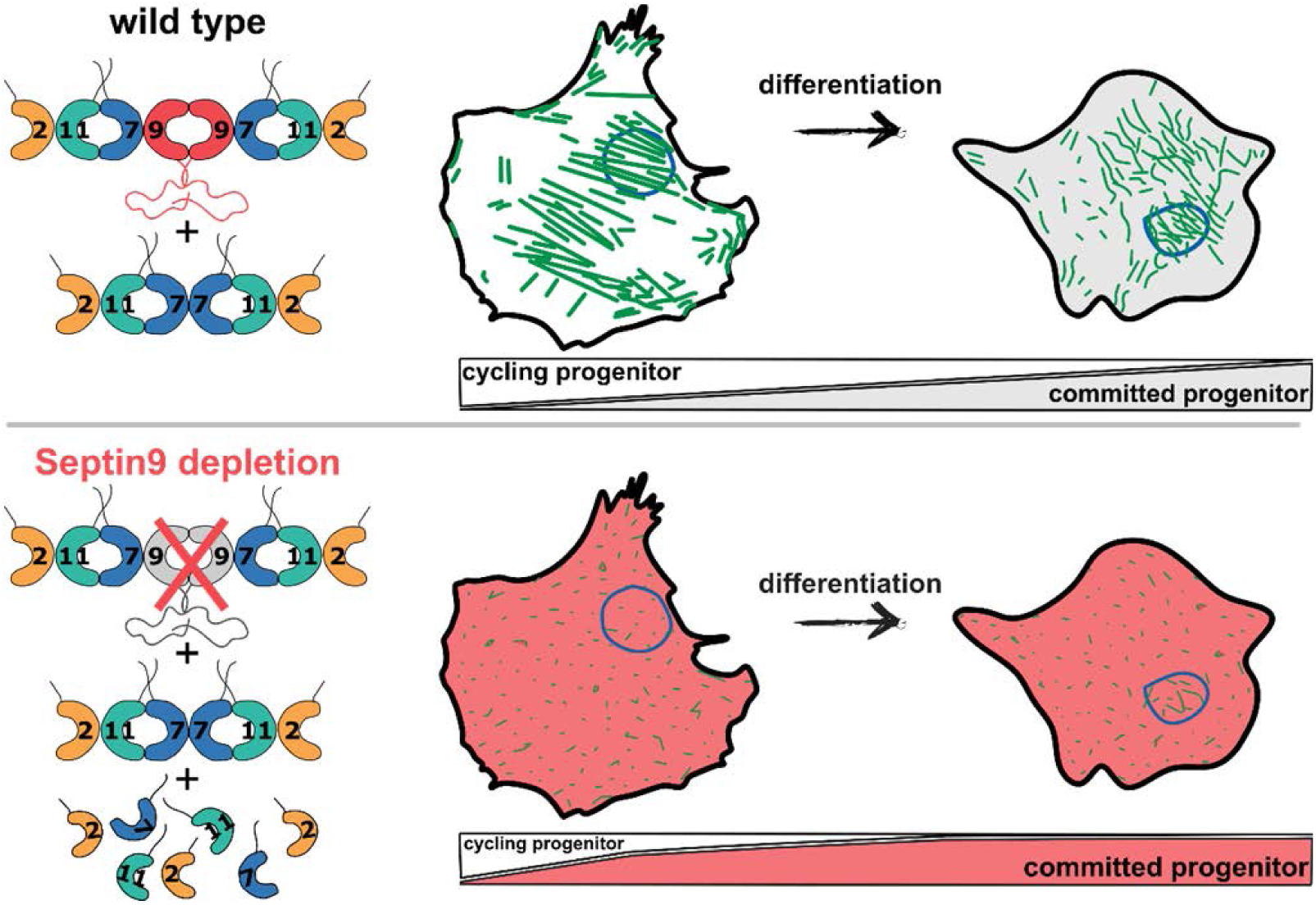
Graphical abstract. The model summarizing our working hypothesis of a premature switch towards committed progenitor program in Septin9 deficient myoblasts. Polymerized, straight, actin-based septin oligomers mark proliferating myoblasts and are strongly present in the perinuclear area. As myoblast transition to migratory myocytes, septins undergo reorganization towards curvy short filaments and presumably become more plasma membrane based. In addition to perinuclear septin-filaments, fusion competent myocytes have largely septin-free protrusions, involved in the initial steps of cell fusion. Septin 9 depletion leads to the collapse of septin filaments in myoblast and premature differentiation towards myocytes

The role of the septin cytoskeleton is poorly understood in the context of cell differentiation. In this study, we use myogenic differentiation as a paradigm and show a requirement of septin structures for regulating the pace of myogenic commitment, and a profound septin filament reorganization during the transition from cycling towards fusion-competent myoblasts. Due to their ability to break symmetry in cells^55^, septins are believed to affect the balance between self-renewal versus differentiation or the mode of stem cell division (symmetric or asymmetric)^56^. In HSCs, Septin7 compartimenatlizes Cdc42 and Borg4, cytoplasmic cell polarity markers, and regulates hematopoietic differentiation potential^33^. Furthermore, by maintaining the Kif20a at the intracellular bridge of NPCs, Septin7 potentially regulates the inheritance of neuronal cell fate determinants by daughter cells. The depletion of Septin7 led to precocious neuronal differentiation, breaking the balanced regulation of asymmetric cell division^32,57^. This raises the possibility that transit-amplifying cells such as myoblasts maintain a limited ability of asymmetric division, or a similar mechanism tied into the balance between proliferation and differentiation. In C2C12-derived myotubes Septin7 was recently shown to interact with Numb, another cell fate determinant^58^. Therefore, it is possible that septin complexes compartmentalize myogenic cell fate determinants via a mechanism resembling neuronal or hematopoietic progenitors and it is conceivable that the role of septins in cell fate determination is conserved across different cell types.One potential limitation of the study is in the difficulty to disentangle the mutually instructing nature of septin-actin interactions. Our *in vitro* data show changes in expression of septin isoforms that are preceded by morphological changes to filament organization, a process that potentially coincides with actin reorganization. Since septins are known to regulate actin stability, bundling and organization, the observed effect of septin depletion might exert its myogenic regulation through impaired actin function^59,60^.

In summary, we show that the septin cytoskeleton is an additional layer of regulation controlling the progression of myogenic differentiation. We propose that the controlled reorganization of septin structures is instrumental in transitioning from an undifferentiated myoblast state to a fusion-competent terminally differentiated state.

## Materials & Methods

### Primary myoblast isolation

Mice were kept in accordance with European Union and German legislation under the license number ZH120. Skeletal muscle progenitors were isolated from hindlimbs of 5 weeks old mice as described in^61^. In brief, muscles were minced and digested with 2.5 mg/ml collagenase A (Roche) for 1 h at 37 °C. Cells were centrifuged at 300 g for 5 min and supernatant was removed. Cell pellet was resuspended in myoblast proliferation medium containing 20% FCS, 10% HS, 2.5 ng/ml recombinant human FGFb (Peprotech), 0.5% chicken embryo extract (LSP). The pellet was placed on Matrigel (Corning) coated plates (working concentration of 0.9 mg/ml) where myoblast migration from the minced myofibers was observed on day 3 of culture. At that point, cells were trypsinized and plated for 1 h on type I rat tail collagen (working concentration 0.1 mg/ml) (Corning) coated plates for removal of fibroblastic and non-myogenic cells. Next, medium containing non-attached cells was collected and placed on Matrigel coated plates for expansion of the myoblast culture.

### Cell culture

C2C12 were obtained from American Type Culture Collection (ATCC) and not used beyond passage 25. C2C12 were cultured in Dulbecco’s Modified Eagle’s Medium (DMEM) containing 1 g/L D-glucose and phenolred (PAN Biotech), supplemented with 10% FCS, 2 mM L-glutamine and penicillin (100 units/mL) / streptomycin (100 µg/mL) (C2C12 full medium) in a humidified atmosphere at 37 °C and 10% CO_2_ (v/v). Primary myoblasts were cultured in DMEM containing 4.5 g/L D-glucose, phenolred and stable L-glutamine, supplemented with20% FCS, 10% HS, 2.5 ng/ml recombinant human bFGF (Peprotech), 0.5% chicken embryo extract (LSP) (myoblast proliferation medium) in a humidified atmosphere at 37 °C and 5% CO_2_ (v/v). To induce differentiation in C2C12 cells, proliferation medium was removed and replaced with differentiation medium: low glucose DMEM, 2% HS (PAN Biotech), 2 mM L-glutamine and penicillin (100 units/mL) / streptomycin (100 µg/mL). To induce differentiation in primary myoblasts, proliferation medium was removed and replaced with differentiation medium: high glucose DMEM, 5% HS and penicillin (100 units/mL) / streptomycin (100 µg/mL).

### Transient transfection with siRNA

Septin9 was silenced in C2C12 using one round of 48h knock-down with 50 nM siRNA, purchased from Dharmacon. Cells were transfected with either scrambled (non-targeting #1) siRNA or Septin9 siRNA (J-048947-11-0050 ONTARGETplus) with Lipofectamine-iMAX (ThermoFisher Scientific) according to the manufacturer’s instructions. In brief, 150.000 cells / 6cm-dish were seeded in 2 ml antibiotic-free medium. On the following day, siRNA – Lipofactamine mix was prepared in Opti-MEM™ – Reduced Serum Medium (ThermoFisher Scientific) and incubated for 20 min. Cells were washed with PBS once and 1.7 ml Opti-MEM™ was added. Next, 300 µl transfection mix was added dropwise to the cells (calculated for total 4 ml transfection medium), effectively increasing the siRNA concentration to 100 nM for the first 5 hours. Subsequently, after 5h 2 ml 20% antibiotic-free medium was added. 24 h later the cells were trypsinized, re-plated for the experiment and the medium was replaced with full proliferation medium. All experiments were performed 48 h after siRNA transfection. Primary myoblasts were handled at recommended densities^61^ and transfected in 3.7 ml full myoblast proliferation medium and 300 µl Opti-MEM/siRNA mix. 24 h later cells were re-plated for the experiment and the medium was exchanged for fresh proliferation medium.

### SDS-PAGE & Western-blotting

For sodium dodecyl sulfate polyacrylamide gel-electrophoresis (SDS-PAGE), treated cells were lysed in 150 µL Laemmli buffer and frozen at –20 °C. The lysate was pulled through a 20-gauge syringe and boiled for 5 min at 95 °C. 10% polyacrylamide gels were cast in advance and stored at 4 °C until usage. Separated by their molecular weight, proteins were transferred onto nitrocellulose membranes by Western blot. Membranes were blocked for 1 hour in 0.1% TBS-T containing 5% w/v skim milk, washed three times in 0.1% TBS-T and incubated with primary antibodies overnight at 4°C. Primary antibodies were applied in 3% w/v bovine serum albumin (BSA)/fraction V in TBST. For HRP-based detection, goat-α-mouse or goat-α-rabbit IgG HRP conjugates (± 0.8 mg/ml) were used in 3% TBST. Chemiluminescent reactions were processed using WesternBright Quantum HRP substrate (Advansta) and documented on a FUSION FX7 digital imaging system.

### Antibodies used in this study

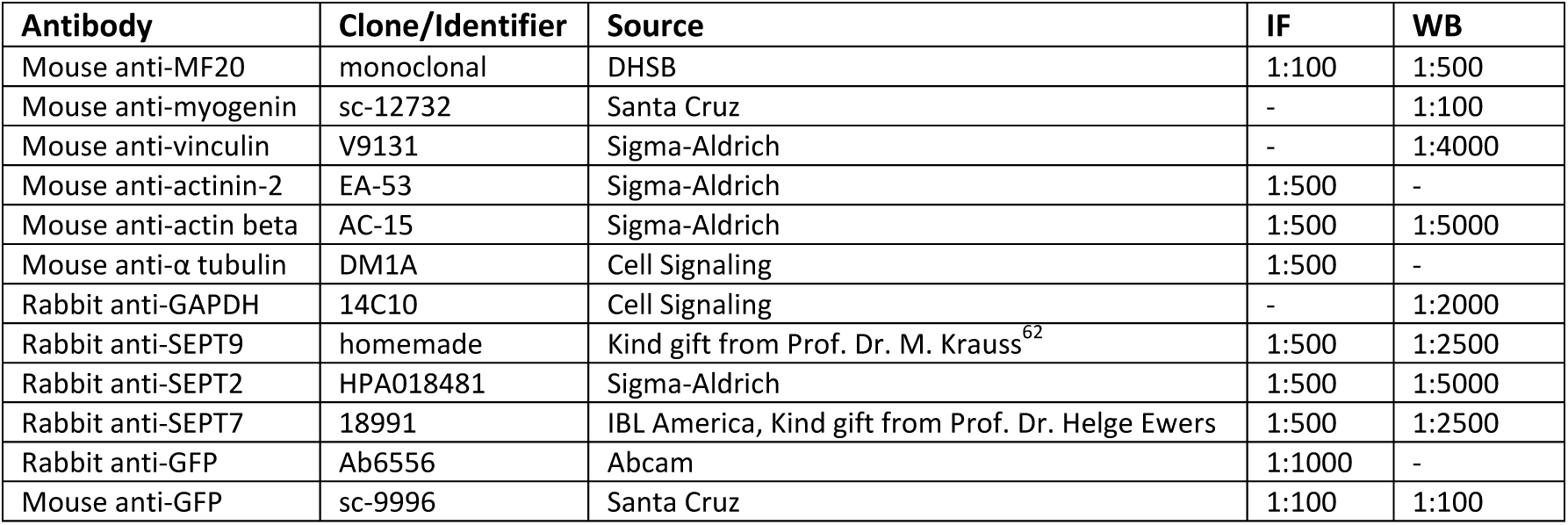
Primary antibodies.

Secondary antibodies.

Secondary antibodies
| Antibody | Conjugate | Source | Identifier | IF | WB |
| --- | --- | --- | --- | --- | --- |
| Goat anti-rabbit | HRP | Dianova | 115-035-068 |  | 1:10000 |
| Goat anti-mouse | HRP | Dianova | 111-035-144 |  | 1:10000 |
| Phalloidin | CruzFluor594 or 647 | Santa Cruz | sc363795, sc363797 | 1:400 |  |
| Goat anti-rabbit | AF 488, 594 | Invitrogen | A11034, A11012 | 1:400 |  |
| Goat anti-mouse | AF 488, 594 | Invitrogen | A11001, A11005 | 1:400 |  |
| Goat anti-rabbit | Atto 647N | Sigma-Aldrich | 40839 | 1:400 |  |

### Generation of a eGFP-Septin9 knock-in cell line

Endogenous tagging of the Septin9 C-terminus with meGFP was achieved via CRISPR/Cas9 ^63^. In brief, the primers 5’-AAACCTGGACCCCACCCCCAGATC-3’ and 5’-CACCGATCTGGGGGTGGGGTCCAG-3’ were annealed and cloned in the px458-pSpCas9(BB)-2A-GFP (Addgene, #48138) using the BpiI restriction site to generate a guide RNA. In a second (“donor”) vector the expression cassette was exchanged with the coding sequence (CDS) of meGFP inserted between two homology regions (HR) consisting of original genomic sequences ∼ 1000 bp upstream (5’HA) and ∼1000 bp downstream (3’HA) of the Septin9 stop codon, separated by a Gly-Ser-Gly-Ser-Gly linker (L). Stop codon of the Septin9 was removed on the donor vector. Design and cloning of the donor vector was performed with the NEBuilder assembly tool and NEBuilder HiFi DNA assembly cloning kit, respectively, according to the manufacturer instructions. C2C12 cells were transfected with px458-pSpCas9(BB)-2A-GFP and donor vector. 72h later GFP-expressing C2C12 cells were sorted into 96-well plates at the density of 1 cell per well, using a fluorescence-activated single cell sorter (BD FACSAriaII SORP (BD Biosciences) at MPI-MG Berlin). Growing colonies were expanded and tested for the expression of Septin9-eGFP by immunofluorescence and Western blotting using anti-Septin9 and anti-GFP antibodies. The expression of Septin9-eGFP in selected clones was further validated by siRNA-mediated depletion of Septin9 and immunocytochemistry.

### Immunocytochemistry

C2C12 cells seeded on 18 µ Ibidi chambers or 12 mm cover slips were fixed with 4% PFA in PBS for 15 minutes at RT and washed two times with PBS. Permeabilization with 0.5% Triton X-100 in PBS was done for 20 minutes, subsequently cells were blocked with blocking solution (5% normal goat serum, 3% BSA in PBS) for 1 hour. Primary antibodies were incubated in the blocking solution for 1 h at RT or overnight at 4 °C, excess of antibody was removed with three washing steps of 5 minutes with 3% BSA in PBS. Incubation with secondary antibodies was carried out in blocking solution for 1 h at RT with three subsequent washing steps. Finally, cells were incubated for 5 minutes with 1 µg/mL of DAPI in PBS and washed with water. 18 µ Ibidi chambers were left in PBS, coverslips were mounted on microscope glass slides with FluoromountG (Invitrogen) or Prolonged Gold (for STED microscopy).

### Confocal & STED microscopy

Confocal and STED data of fixed C2C12 cells were acquired with the Expert Line STED Microscope from Abberior. Confocal images of Septin9-GFP C2C12 cells were acquired using 485 nm (20% laser power) and 640 nm excitation (20% laser power). STED images were acquired using 561 nm excitation for ATTO 594 (20% laser power) and 640 nm excitation for a 775 nm STED laser at 10% laser power was used to deplete both dyes.

### Live cell microscopy

Live cell imaging was performed in differentiation medium without phenolred at 37 °C and 5% CO2. Confocal live cell imaging of Septin9-eGFP cells was performed on a spinning disk Nikon Eclipse Ti microscope (Yokogawa CSU-X1 and EMCCD Camera), operated by NIS-Elements software, with a 40× air objective (0.75 NA). Imaging was initiated 72 h after medium change, and was carried overnight, with a frame rate of 30 minutes. Pictures were acquired within two z-planes that were set at the beginning by focusing on septin fibers and 0.5 µm above, and kept by an autofocus system. TIRF live cell imaging was conducted with a Nikon Eclipse Ti microscope (illumination: TIRF laser 488, prime95B sCMOS camera) operated by Micromanager, using a 60× oil objective (1.49 NA). Imaging was initiated after medium change, and was carried overnight, with a frame rate of 30 minutes

### Image analysis & semi-automated quantification with Fiji

For figure 3 at least 3 independent experiments with 3 technical replicates from the 8 µ Ibidi chamber slide were performed. Tile scans of 6.4 (4×4) or 3.6 (3×3) μm^2^ were produced by stitching together images automatically acquired around a chosen point with 20× objective. Four to five tile scans were acquired per condition. The progress of myogenesis was quantified by counting nuclei within and outside of myotubes using a custom-written Fiji macro. Both myotubes and nuclei were fluorescently labeled (MF20 and DAPI), and a Gaussian blur with a sigma radius of 1 pixel was applied to smooth the color channels. The Otsu-based thresholding method was employed to separate the nuclei from the background, and a watershed algorithm was utilized to segment touching nuclei. Morphological operations including despeckle, 4× dilate, close, fill holes, and 3× erode were performed on the stained myotubes to address any irregularities. Additionally, touching myotubes were segmented using a watershed algorithm. The processed myotube channel was used to define regions of interest (ROIs), and the number of nuclei was counted for each myotube within these defined regions. To quantify the total number of nuclei, both within and outside of the myotubes, Fiji’s Particle Analyzer was employed. To ensure the accuracy of the analysis, a composite image combining the fluorescence channels of the myotubes and nuclei was generated. The outlines of the analyzed myotubes and nuclei were overlaid on this image, allowing for manual validation of the analysis quality.

### Quantitative real-time PCR

Myogenic differentiation experiments with C2C12 and primary myoblasts was initiated by a medium change for serum-reduced differentiation medium after cells reached 90% confluency. For sample collection, cells were rinsed twice with PBS and lysed in 350 µl RA1 buffer (RNA extraction kit), supplemented with 1% β-mercaptoethanol, stored at –80 °C until further processing. Cellular RNA was isolated using the NucleoSpin RNA isolation kit (Macherey-Nagel, Düren, Germany) according to the manufacturer’s instructions. 1 μg total RNA was reversely transcribed by incubation with random primers (100 pmol μL^−1^, Invitrogen, Carlsbad, USA) and M-MuLV reverse transcriptase enzyme (200,000 U mL^−1^, New England Biolabs, Ispwich, USA) was added per sample. RT-PCR was performed using a StepOnePlus Real-Time PCR System (Thermo Fisher Scientific) with specific primers. Reactions were performed in triplicates in MicroAmp Optical 96-well reaction plates (Thermo Fisher Scientific) using Luna PCR Master Mix (New England Biolabs). Fold induction was calculated by comparing relative gene expression to the housekeeping gene 18S RNA using the ΔΔCT method^64^.

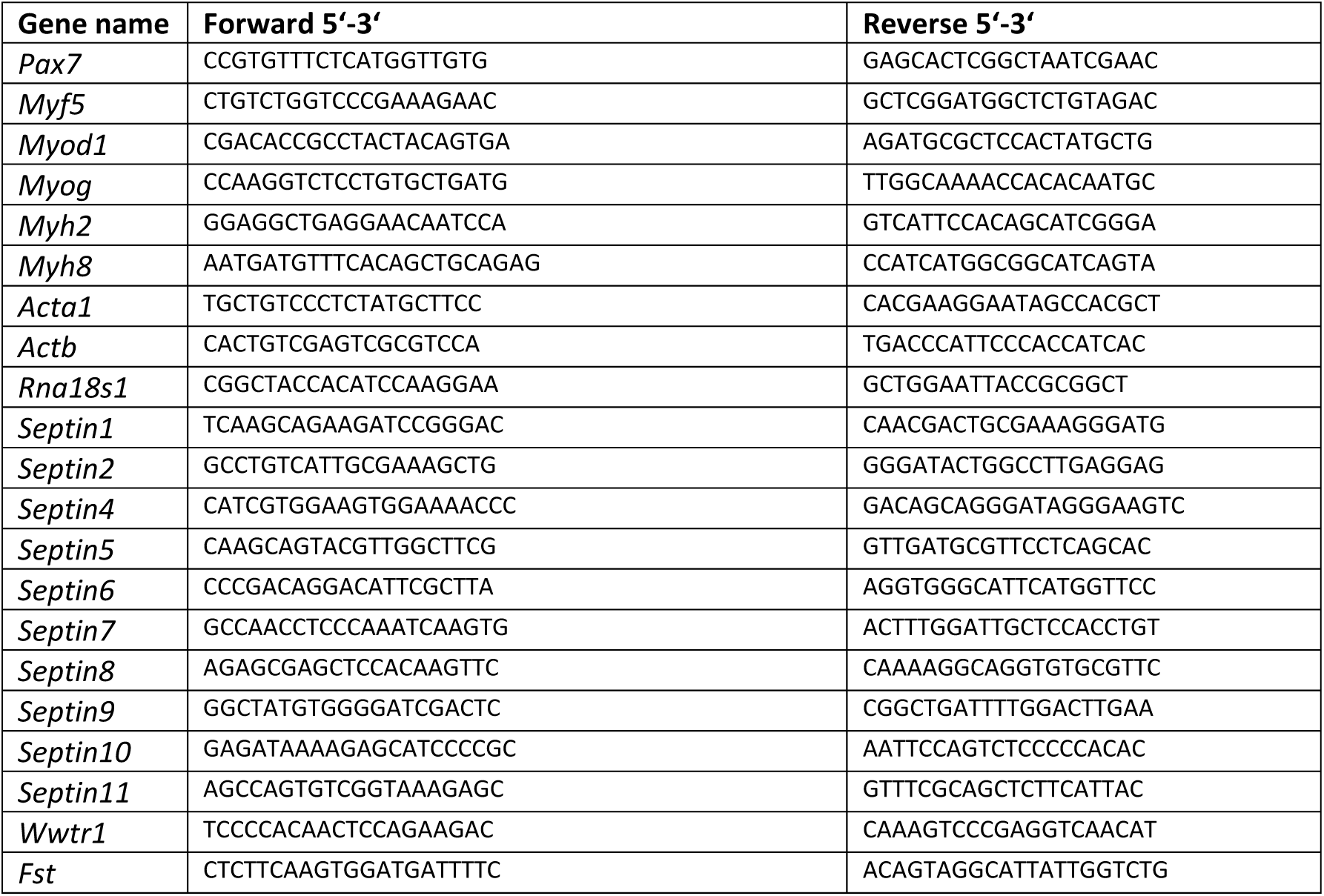
Primers used for qRT-PCR.

### Single-nucleus RNA-sequencing data analysis

Single-nucleus RNA sequencing data was taken from the mouse organogenesis cell atlas (MOCA)^34^. The sparse count matrix was downloaded from the MOCA data portal (https://oncoscape.v3.sttrcancer.org/atlas.gs.washington.edu.mouse.rna/downloads). The initial matrix was subset to only include cells annotated by the authors as mesenchymal lineage. The raw count matrix was processed using Seurat v3^65^. Data was first log-normalized using a scale factor of 10^4^. The top 5000 variable genes were identified by the *FindVariableGenes()* function using the default settings. Centering and scaling of the data using a linear regression model was then performed on the previously identified 5000 variable features, regressing out the number of unique molecular identifiers (UMI) and the percentage of mitochondrial reads per cell. For clustering, a shared nearest neighbor graph was constructed using the first 30 principal components (PCs). Louvain clustering was performed with the clustering resolution set to 0.8. Clusters were annotated based on expressed marker genes. The differentially expressed marker genes upon which cell identities were assigned are provided below. Clusters annotated as progenitors, myoblasts and myocytes were defined as the myogenic lineage. The myogenic lineage was then extracted and exported as raw counts.

### Pseudotime analysis

For trajectory inference, the raw count matrix of the myogenic lineage was analyzed using the default parameters of Monocle 3^34^. Briefly, cells were log-normalized and scaled followed by principal component analysis. After Louvain clustering, dimensionality was reduced by uniform manifold approximation and projection (UMAP)^66^ initialized on the first 30 PCs. Finally, the principal graph was fitted, where progenitor cells were assigned as the root of the trajectory.

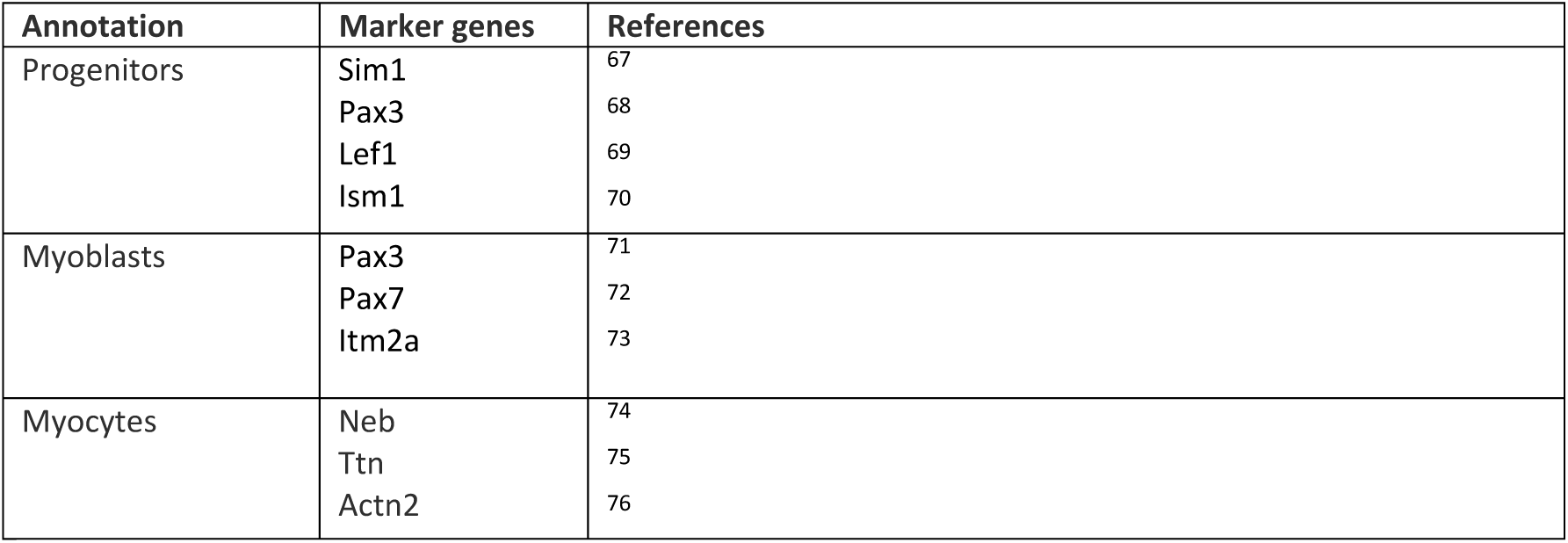
Cell annotation.

### RNA-seq library preparation and sequencing

40.000 cells/cm² cells were seeded in a 12 well plate 24 hours after Septin9 knockdown in full myoblast proliferation medium from two independent experiments. Cells were expanded for 24 hours (total 48 hours after knockdown). Primary myoblasts were lysed, and C2C12 cells differentiated for 12 hours and then lysed. RNA was isolated according to the manufacturer instructions (NucleoSpin RNA isolation kit (Macherey-Nagel)). 500ng RNA was sent for sequencing to Genewiz, Leipzig, Germany.

### RNA-seq data analysis

RNA-seq analysis was performed using the Galaxy platform^77^. Paired-end, 300-bp reads from Illumina sequencing were mapped to the GRCm38.p6 reference genome with STAR (v2.7.2b)^78^ and default settings. Aligned reads were assembled using StringTie (v2.1.1)^79^. Subsequently, batch effects in the resulting count data were detected with RUVSeq^80^ (k=1) and gene expression was quantified using DESeq2^81^ on wild-type versus knock-down samples with batch effects removed by including the RUVSeq results. Gene ontology enrichment analysis was performed with Enrichr^82^. Subsequent data visualization was performed in R (v4.1.1) using the pheatmap^83^, ggplot2^84^, ggrepel^85^, and dplyr^86^ packages.

### Statistical analysis

All data are derived from at least three independent experiments (except RNA-seq) and are represented as means± standard deviation (SD). Statistical tests were performed using GraphPad Prism (v9.3) software. All statistical tests are listed in the figure legends. Datasets were tested for normality with the Shapiro-Wilk test. Two tailed student’s t-test was used to compare between two conditions. Whenever comparing more than two conditions, the one-way ANOVA and Dunnett post hoc test were used to check for statistical significance under the normality assumption. The level of significance is indicated in the figures by asterisks (*P < 0.05; **P < 0.01; ***P < 0.001; ****P < 0.0001).

## Acknowledgments

V.U. was supported be the Deutsche Forschungsgemeinschaft DFG (SFB958, SFB1444) and Sonnenfeld Stiftung. PM was supported by the Max-Planck Research School (IMPRS-Biology and Computation). J.J. was supported by the Deutsche Forschungsgemeinschaft DFG (BSRT, SFB958) and the Einstein Center ECRT. P.K. acknowledges the support from Deutsche Forschungsgemeinschaft DFG (FOR2165; SFB1444), the Einstein Center ECRT, Morbus Osler Society, and BMBF (PrevOP-Overload). We are very grateful to the Cellular Imaging Facility of the Leibniz-Forschungsinstitut für Molekulare Pharmakologie (FMP) for access and support with live-cell imaging and the animal facility of the Max Planck Institute for Molecular Genetics for expert support.

## Author contributions

V.U. conducted experiments, P-L.M contributed RNASeq data analysis, D.Z. helped with *in vitro* characterization of C2C12, G.R. helped with knock-in cell line generation and live cell microscopy, Y.K. developed a script for automated myogenic fusion index quantification, T.H. contributed snRNASeq data analysis, G.K. helped with primary myoblast isolation and maintenance, J.J. conducted STED imaging, S.S and P.K supervised the study, P.K together with V.U. wrote the paper with input from all authors.

## Figures

**Figure EV1.**
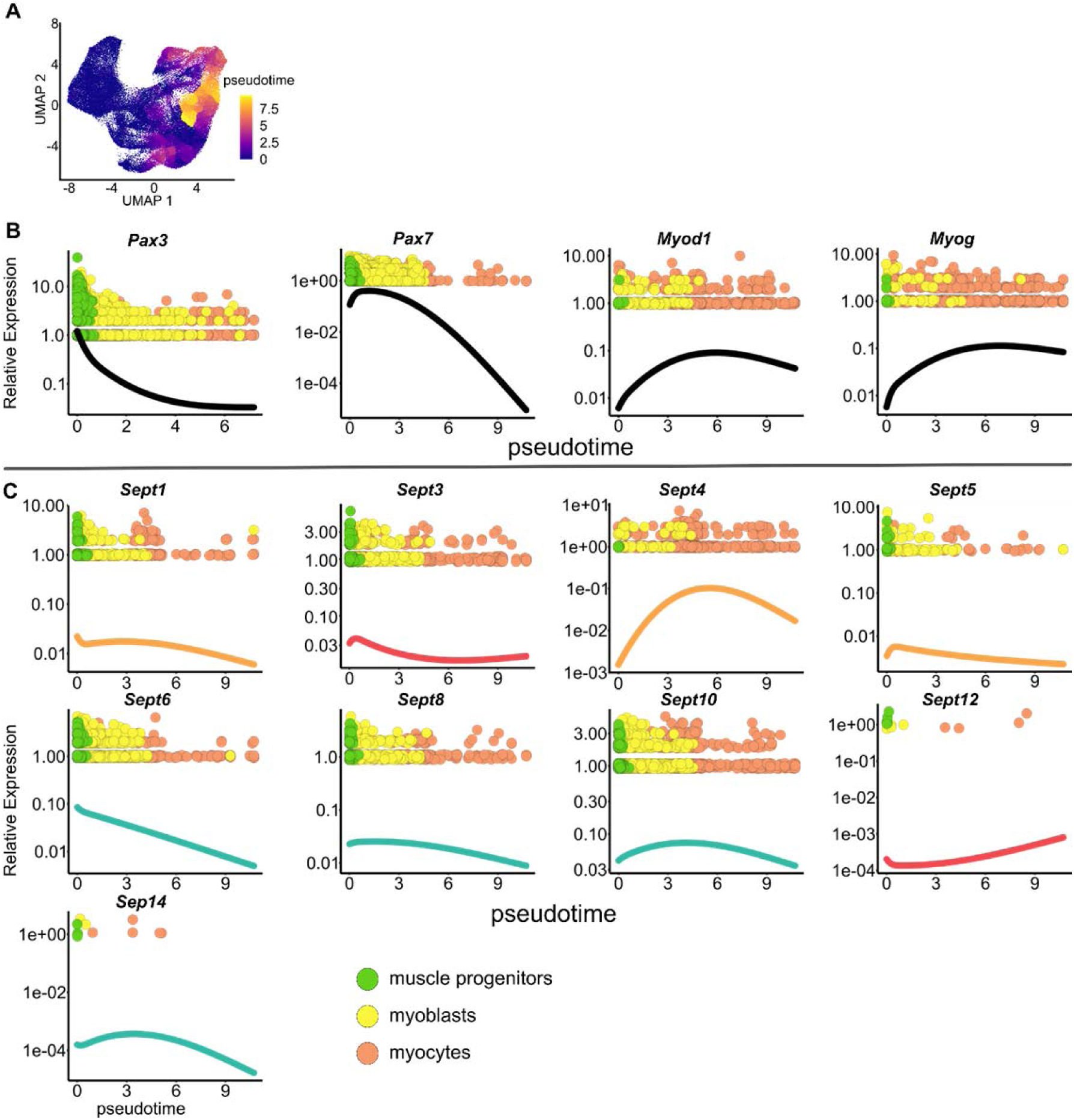
Transcriptional landscape of septins during mouse musculature development. **a** UMAP visualization of the developing musculature ranging from embryonic day 9.5 to 13.5, colored by inferred pseudotime. Kinetics plots showing relative expression of **b** myogenic marker genes and **c** all septin paralogs across developmental pseudotime. Spline curves are color coded by protomer structure proposed in Fig.1d.

**Figure EV2.**
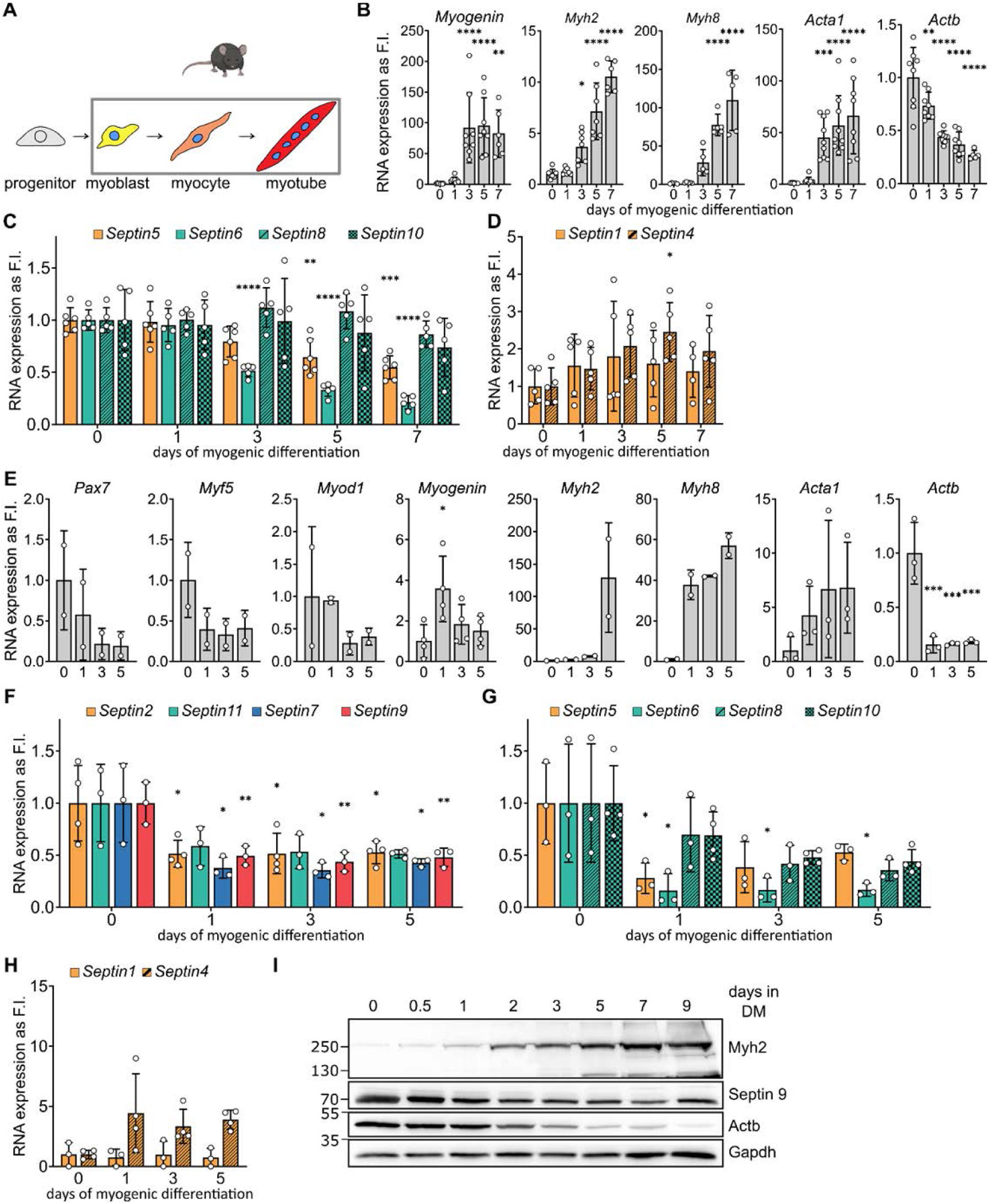
Expression of septins during myogenesis in C2C12 cells and primary myoblasts. **a** Schematic representation of murine *in vitro* myogenesis. **b** mRNA expression of myogenic marker genes during 7 days of myogenic differentiation of C2C12 cells. **c-d** mRNA levels of other expressed septin paralogs. **e** mRNA expression of myogenic marker genes during 5 days of myogenic differentiation of primary myoblasts **f** mRNA expression levels of core myogenic septins and **g-h** other expressed septin subunits. **i** Representative western blot of Septin9 during 9 days of myogenesis in primary myoblasts. Data represent mean ± standard deviation (SD), *p<0.05, **p<0.01, ***p<0.001, ****p<0.0001 from one way ANOVA followed by Dunnett’s multiple comparison test.

**Figure EV3.**
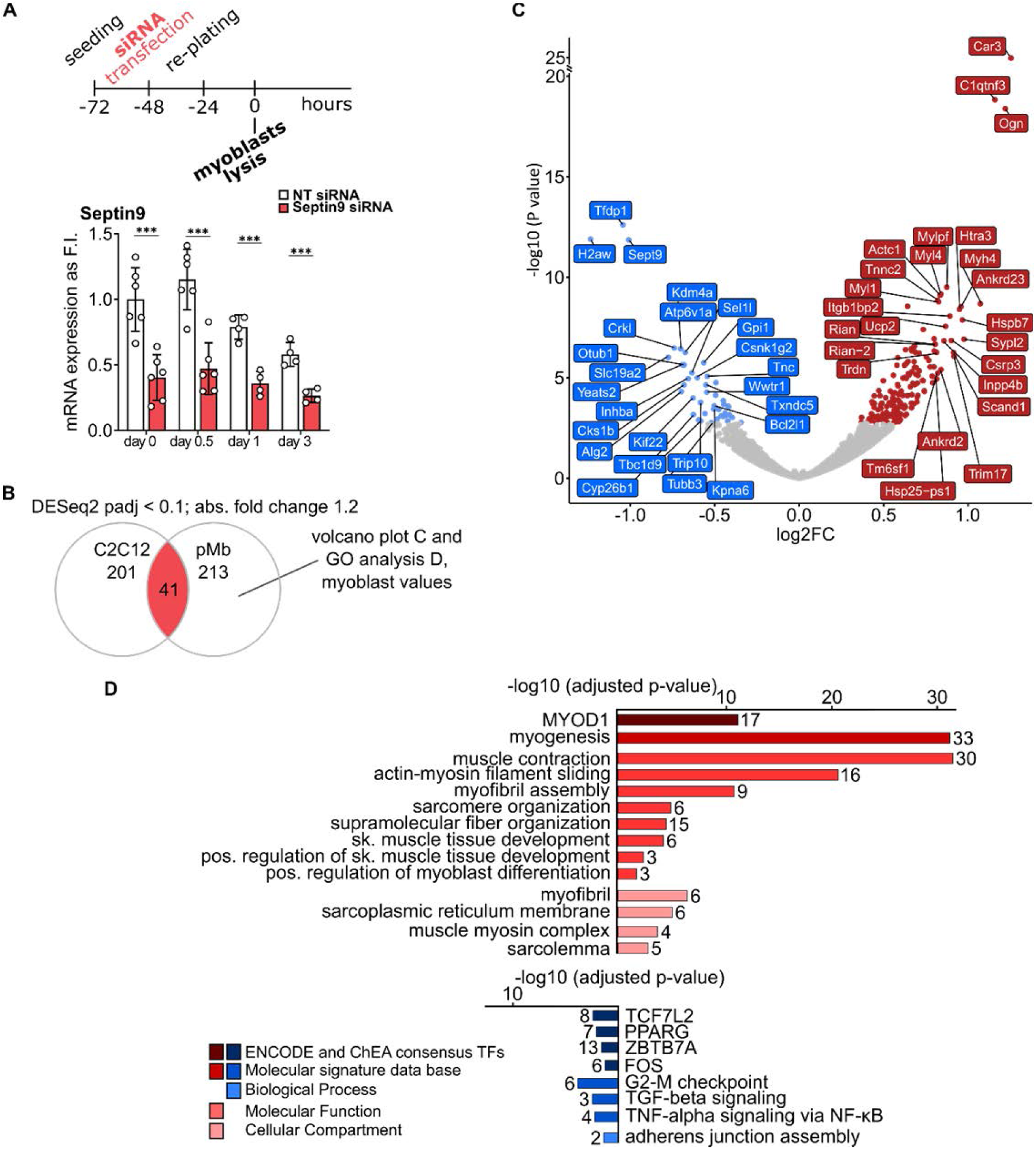
Septin9 depletion accelerates myogenic differentiation in primary myoblasts. **a** Schematic representation of the experimental setup. Total RNA was isolated from proliferating myoblasts 48 hours after Septin9 knockdown. Septin9 knockdown efficiency validation over 3 days of in vitro myogenesis via qRT-PCR in primary myoblasts. **b** Venn diagram comparing DEGs calculated using Dseq2 algorithm (adjusted p value < 0.1; –0,322 ≤ log2FC ≤ 0,263). **c** Volcano plot of DE genes (adjusted p value <0.1; –0,322 ≤ log2FC ≤ 0,263) in primary myoblasts transfected with either non-targeting siRNA (control) or Septin9 siRNA, 50 most regulated DEGs are highlighted. **d** GO enrichment analysis of genes upregulated (red) or downregulated (blue) in Septin9-deficient myoblasts. Data represent mean ± standard deviation (SD), *p<0.05, **p<0.01, ***p<0.001, from two-sided unpaired t test.

**Figure EV4.**
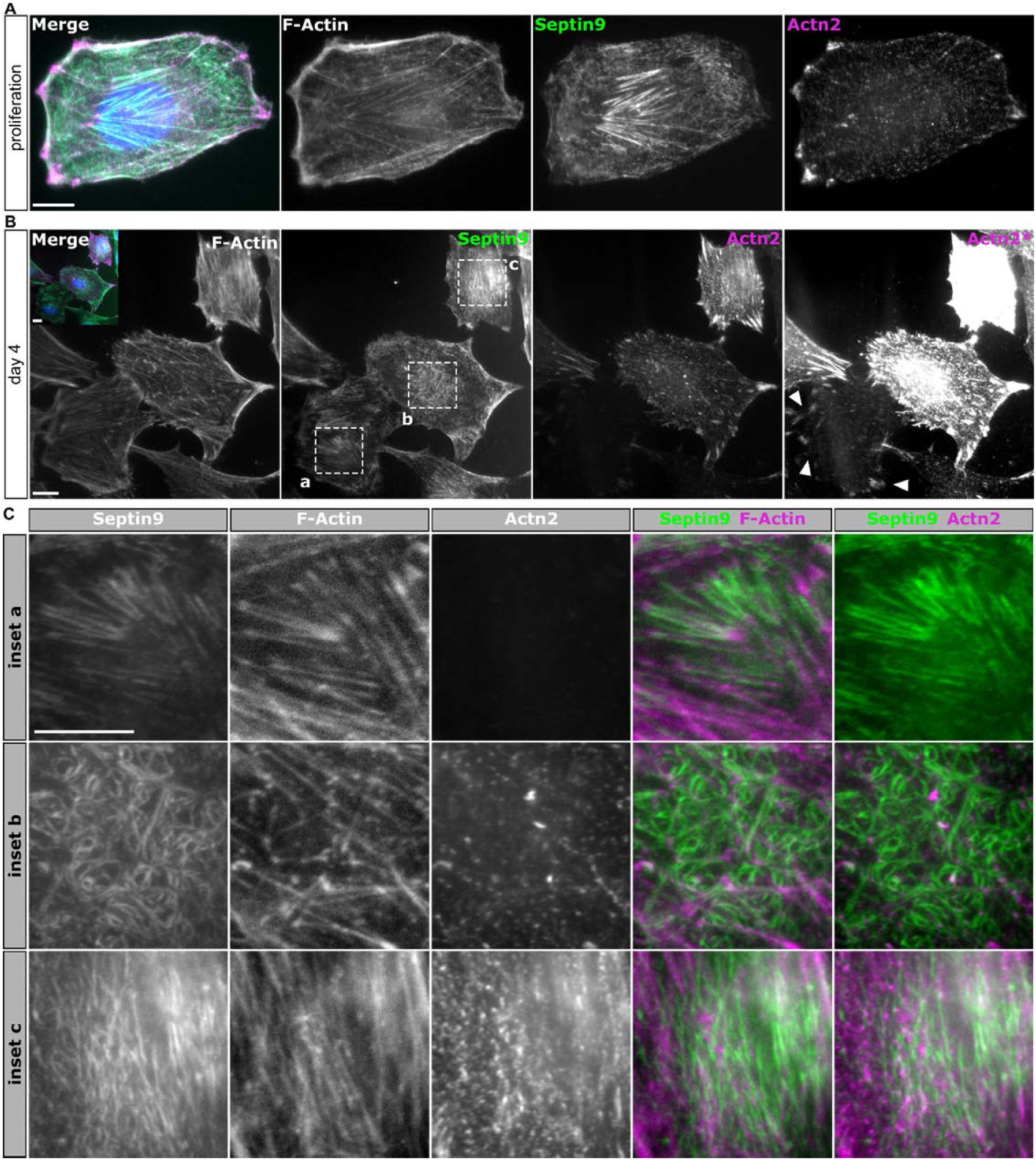
Endogenous Septin9 does not colocalize with α-actinin-2 during myogenesis. **a** Representative epifluorescence images of proliferating WT C2C12 cells, visualizing α−actinin-2, Septin9 and F-Actin via phalloidin. **b** Representative epifluorescence images of mononuclear myoblasts differentiating for 4 days. Undifferentiated myoblasts express low levels of α-actinin-2, localized to focal adhesions (arrow heads). Committed myoblasts increase α−actinin-2 expression in preparation for sarcomere assembly. Septin9 dissociates from actin and does not colocalize with α−actinin-2 during this transition. The contrast in the “α-Actn2 channel” was modified (marked as Actn2*) for better visibility of the cell with the low α-actinin-2 level. **c** Magnified insets from **b**, emphasizing Septin9 reorganization and the absence of Septin9-α−actinin-2 colocalization in differentiating myoblasts. Scale bar 10 µm, inset 5 µm.

**Figure EV5.**
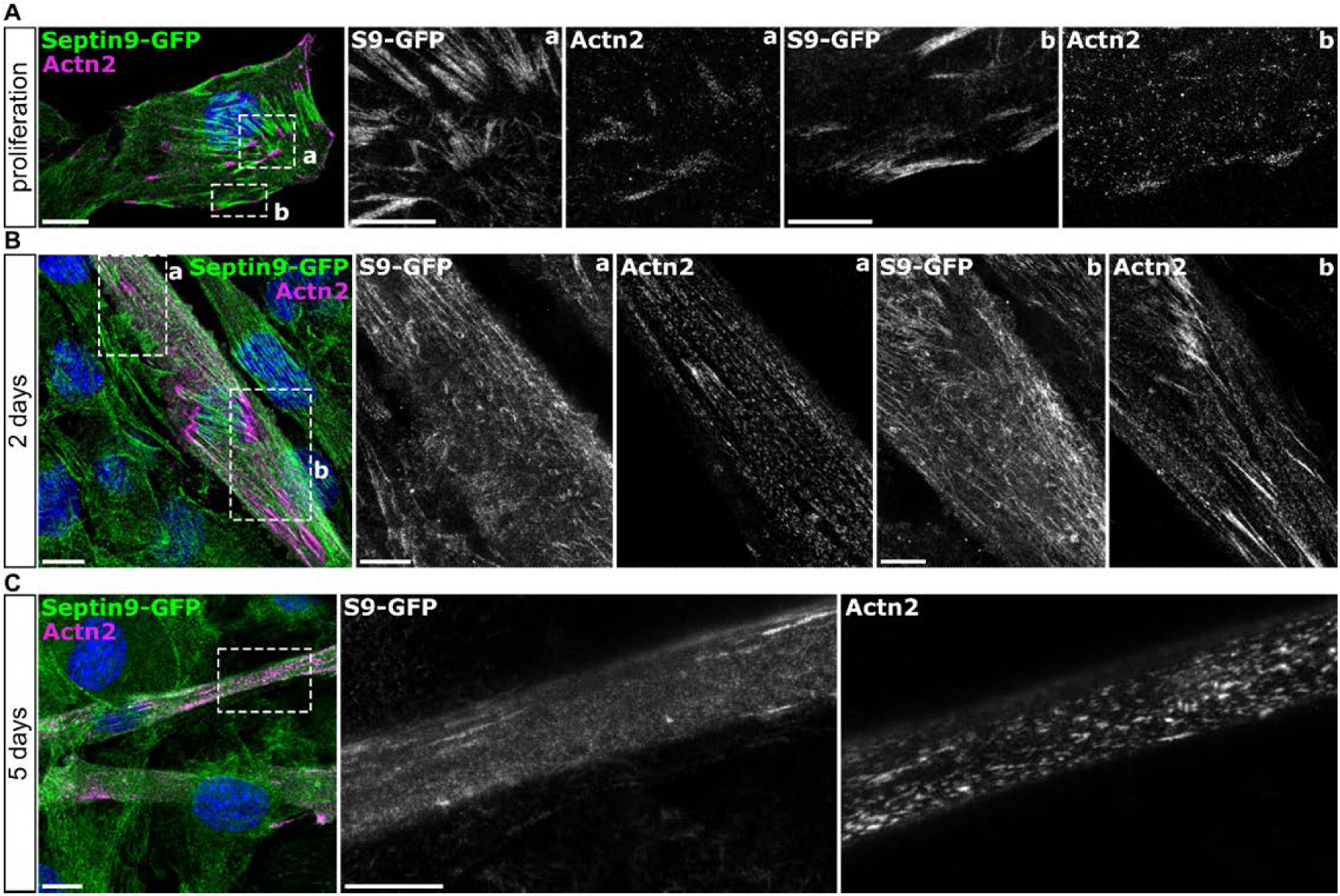
Septin9-GFP does not colocalize with α-actinin-2 during *in vitro* myogenesis. **a** Confocal (merge) and STED (insets) fluorescent micrographs of proliferating Septin9-GFP knock-in myoblasts. α-actinin-2 localizes to the focal adhesions and Septin9-GFP is excluded from these structures. **b** Representative images capture cytoskeletal reorganization in the nascent myotube after 2 days of differentiation. Septin9-GFP is no longer organized in long, straight filaments, but in short rods and rings, showing no colocalization with α-actinin-2. **c** The mature myotube shows no colocalization between residual Septin9-GFP and α-actinin-2, that resembles primitive sarcomeres. Scale bars 10 µm, insets in cells 5 µm.

**Figure EV6.**
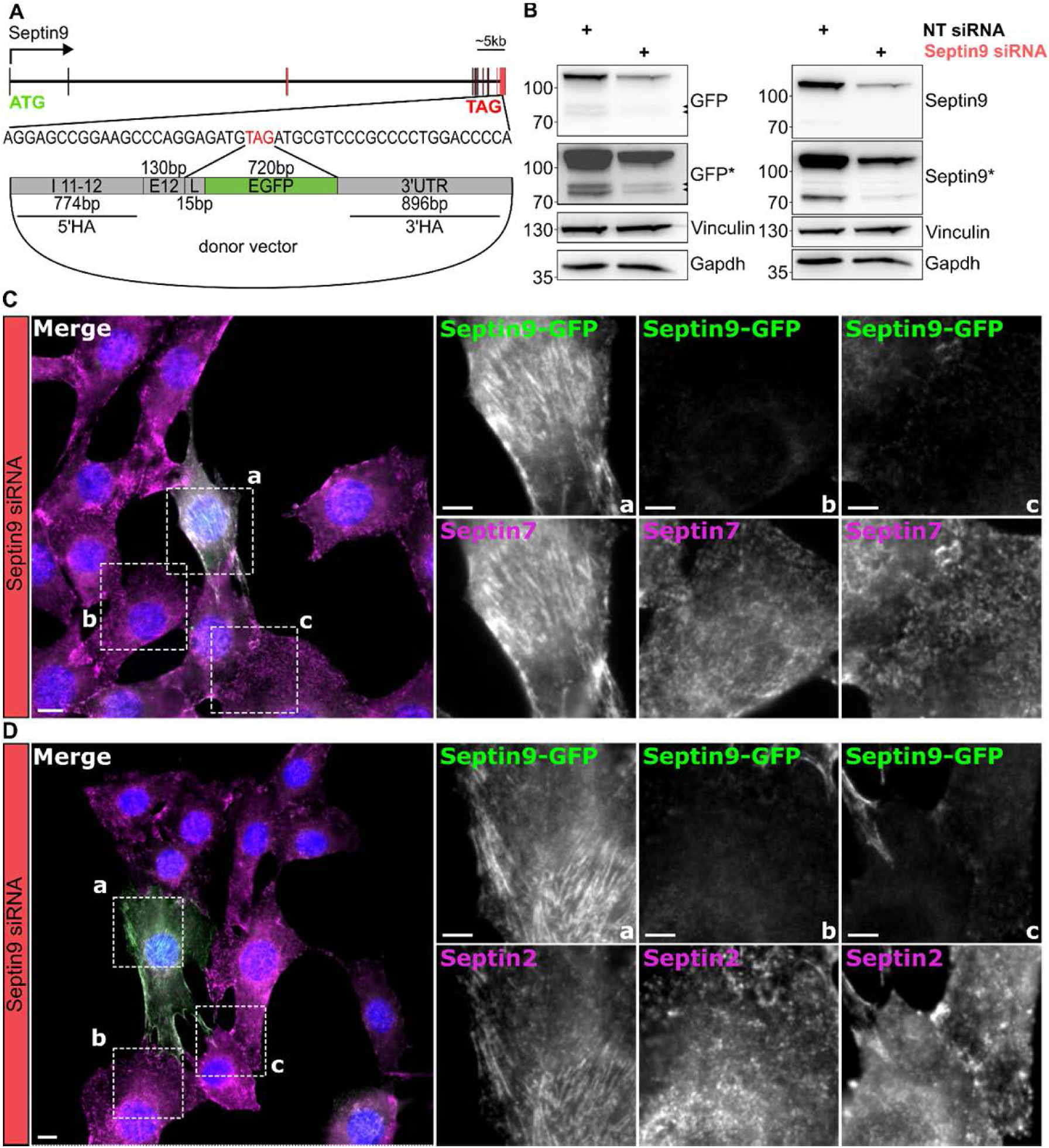
Generation and validation of a Septin9-GFP C2C12 cell line. **a** Cloning strategy introducing a 15bp linker (L) and an EGFP sequence instead of the stop codon of Septin9 in C2C12 cells. C-terminal strategy targets all Septin9 isoforms. Several clones were generated and tested. **b** Clone 32 was transfected with either non-targeting siRNA or Septin9 siRNA and subjected to a SDS PAGE analysis followed by a western blot. Antibodies against long Septin9 isoforms (run above 70kDa) and GFP were used to validate the knock-in. The molecular weight shift of about 30kDa (GFP) is visible with both antibodies. Probing the membrane with anti-GFP antibody might reveal the expression of the short isoforms (black arrow heads). Lanes marked with an asterisk (GFP* and Septin9*) represent longer exposure. **c-d** Representative epifluorescent images of proliferating Septin9-GFP clone#32 48 hours after siSeptin9 treatment visualizing Septin7 (**c**) and Septin2 (**d**). Sepin9-GFP shows knock-in GFP signal, Septin7 and Septin2 are stained with antibodies. Scale bar 10 µm, insets 5 µm.

**Figure EV7.**
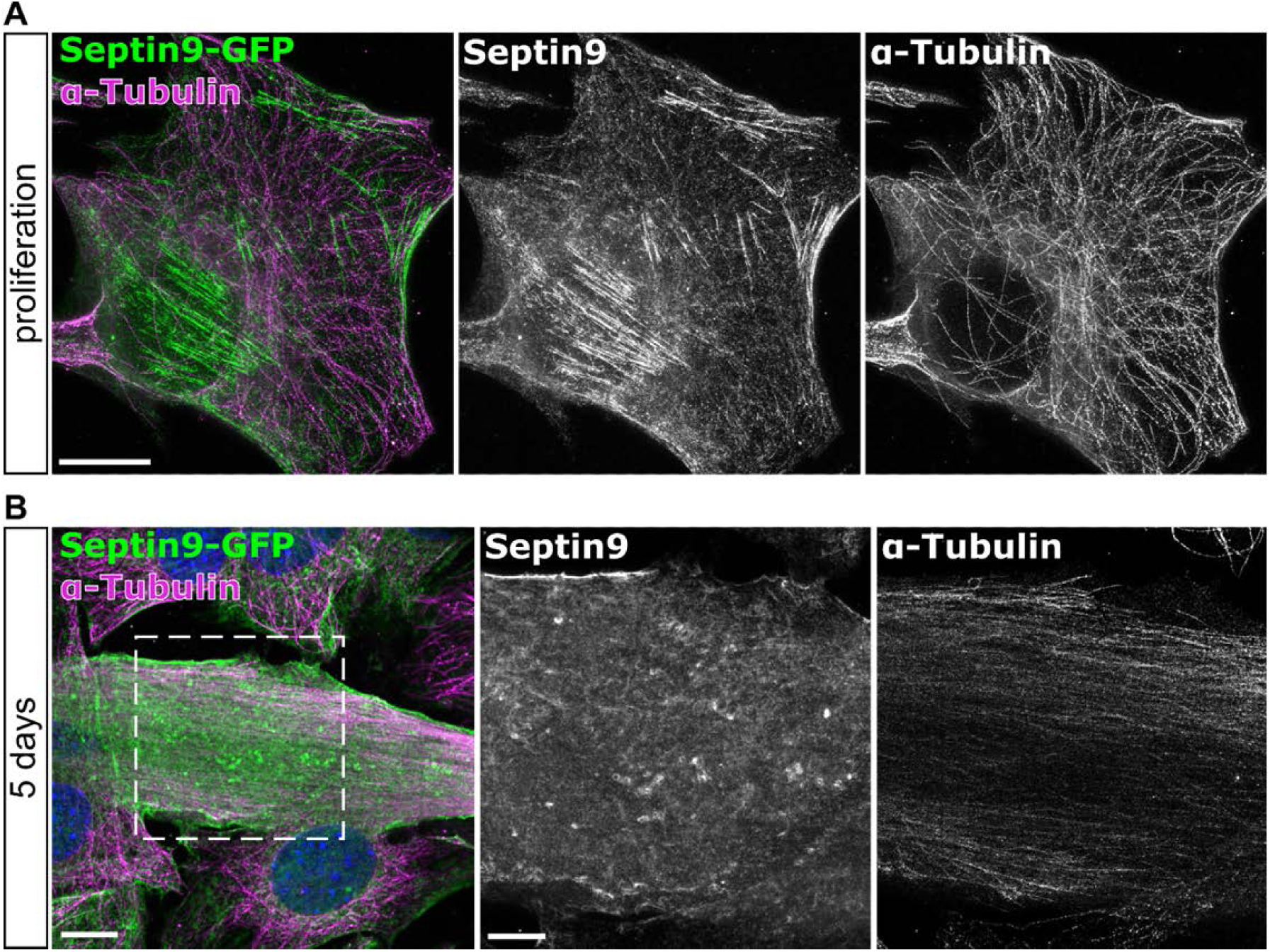
Microtubules in C2C12 cells are mostly devoid of Septin9. **a** Confocal (merge) and STED (insets) fluorescent micrographs of proliferating and differentiating for 5 days (**b**) Septin9-GFP knock-in myoblasts. Microtubules are visualized using α-Tubulin antibody. Septin9 appears to colocalize only occasionally at the cell periphery. Scale bars 10 µm, insets in cells 5 µm.

